# Selenium Silk Nanostructured Films for Antifungal and Antibacterial Treatments

**DOI:** 10.1101/2022.12.18.520325

**Authors:** Zenon Toprakcioglu, Elizabeth G. Wiita, Akhila K. Jayaram, Rebecca C. Gregory, Tuomas P. J. Knowles

**Affiliations:** Yusuf Hamied Department of Chemistry, University of Cambridge, Lensfield Road, Cambridge CB2 1EW, UK; Cavendish Laboratory, Department of Physics, University of Cambridge, J J Thomson Avenue, Cambridge CB3 0HE, UK

**Author notes:** These authors contributed equally (Zenon Toprakcioglu, Elizabeth G. Wiita, Akhila K. Jayaram).

## Abstract

The rapid emergence of drug-resistant bacteria and fungi poses a threat for healthcare worldwide. The development of novel effective small molecule therapeutic strategies in this space has remained challenging. An orthogonal approach, therefore, is to explore biomaterials with physical modes of action which have the potential to generate antimicrobial activity and in some cases even prevent antimicrobial resistance. Here, to this effect, we describe an approach for forming silk-based films that contain embedded selenium nanoparticles. We show that these materials exhibit both antibacterial and antifungal properties while crucially also remaining highly biocompatible and non-cytotoxic towards mammalian cells. By incorporating the nanoparticles into silk films, the protein scaffold acts in a two-fold manner; it protects the mammalian cells from the cytotoxic effects of the bare nanoparticles, while also providing a template for bacterial and fungal eradication. A range of hybrid inorganic/organic films were produced and an optimum concentration was found, which allowed for both high bacterial and fungal death while also exhibiting low mammalian cell cytotoxicity. Such films can thus pave the way for next generation antimicrobial materials for applications such as wound healing and as agents against topical infections, with the added benefit that bacteria and fungi are unlikely to develop antimicrobial resistance to these hybrid materials.

## Introduction

Biomaterials derived from natural sources, including proteins and peptides, provide a unique opportunity to create biocompatible structures for biomedical applications.^1^ The self-assembly of these protein molecules can lead to functional complexes that have tunable three-dimensional structures, ranging in size from nanometers to centimeters.^2–5^ Furthermore, these structures offer promising routes for loading, carrying, and releasing cargo molecules to selected targets. More importantly, protein-based structures are nontoxic, non-immunogenic, biodegradable, and biocompatible, making them ideal candidates for drug/gene delivery applications.^6–10^ A group of materials that distinctly match these applications due to their bio-based components are silk-derived proteins. ^2,8,11–13^ In its natural role, native silk fibroin (NSF) is spun into long fibers by the *Bombyx mori* silkworm, resulting in strong hydrogen-bonding within *β*-sheet nano-crystallites that have both highly ordered and amorphous regions, giving the silk threads their durable mechanical properties.^14,15^ NSF can be dissolved to form regenerated silk fibroin (RSF),;^14,16–18^ resulting in a new material that is a natural block copolymer, which can be readily reassembled into bespoke structures that retain many of the attractive chemical and physical properties that native silk possesses, including durability, biocompatibility, and tunability.^19^ These qualities make it an excellent resource in medical materials, tissue engineering, and in the delivery of therapeutics, with the advantage that RSF can be handled with greater ease than its native counterpart. ^11^ Recently, RSF has been used in several medical applications including forming antibacterial materials, wound healing matrices, and targeted drug delivery gels.^2,7,12,20^ Moreover, there has been an increased interest in developing RSF based structures which are integrated with nanoparticles, such as selenium nanoparticles, in order to produce antimicrobial materials^21,22^ that are more prone to killing drug-resistant bacteria and fungi.

Antimicrobial resistance remains one of the most critical public health concerns, as current medicines used to treat a variety of ailments caused by viruses, fungi, and bacteria become ineffective.^23–25^ Research into antimicrobial agents that subsequently do not become resistant to such microbes is thus needed to avert this crisis. Several studies have focused on silver nanoparticles for antimicrobial solutions; however, studies have indicated emerging resistance to these materials from bacterial populations as well as toxicity to human cells.^26–28^ One potential alternative to silver in this role is selenium, a trace element in the human body that is an essential micronutrient often found as selenocysteine, the 21st human amino acid.^29,30^ Selenium nanoparticles have been shown to exhibit effective antimicrobial properties^31–33^ against a broad range of microorganisms including *Escherichia coli* and *Candida albicans*.^32^ Moreover, there has been an increased interest in functionalising selenium nanoparticles using chitosan,^34^ hyaluronic acid,^35^ polyethylene glycol (PEG),^36^ pectin^37^ and peptides,^38^ in order to improve stability and enhance biocompatibility. ^39^ Seleniums naturally occurring presence in the human body makes it an excellent alternative for antimicrobial development because it is much less toxic to humans than silver and is unlikely to develop antimicrobial resistance. ^40–42^ Importantly, the antimicrobial properties and efficacy of selenium nanoparticles depend on their size and arrangement within a material to effectively inhibit microbial growth.

In this study, we have developed hybrid selenium-silk based films capable of potent antimicrobial properties and with excellent biocompatibility. In order to do this, we first optimised the formation of selenium nanoparticles through the reduction of sodium selenite by ascorbic acid.^43^ By varying the concentration of sodium selenite from 100-1000 μg/mL, we were able to form nanoparticles of different sizes. Using electron microscopy, we determined the size distribution of the nanoparticles over a 6-day period. We found that nanoparticle diameters varied (60-130 nm) as a function of the sodium selenite concentration. Additionally, the stability over time was monitored and it was found that the nanoparticles agglomerated as time progressed, with complete agglomeration happening by day 6. This issue was addressed by combining the nanoparticles with silk fibrils, which acted as stabilisers and did not allow the nanoparticles to clump together. These hybrid silk-selenium films demonstrate unique malleability and have a homogeneous nanoparticle distribution within the protein matrix. Finally, we looked at the potential use of these films as antimicrobial agents. Three different bacteria and fungi were investigated; the gram-negative bacterium *E. coli*, the gram-positive bacterium *Bacillus subtilis* and the fungi *Candida parapsilosis*. In all systems, it was determined that the films displayed potent antifungal and antibacterial properties, killing the majority of the bacteria/fungi. More importantly, however, we found that by incorporating the selenium nanoparticles within the films, mammalian cells were protected against the cytotoxic effects of the bare nanoparticles. Therefore, we were able to not only retain the antimicrobial effect of the nanoparticles, but also ensure that our material was highly biocompatible, making these silk-selenium films ideal for use as topical antimicrobial agents.

## Results and Discussion

In order to generate a material which displayed antimicrobial properties while also being biocompatible, we chose to investigate a hybrid inorganic/organic system. As selenium has been shown to exhibit antimicrobial properties, selenium nanoparticles (SeNPs) were synthesised using a redox reaction. The SeNPs were then integrated with pre-formed silk fibrils, before being left to self-assemble, resulting in the formation of a Silk-SeNP film (the schematic of which is shown in Figure 1). The films were then tested with fungi, gram negative and gram positive bacteria, and it was found that in all cases microbial death was observed. Finally, viability assays were conducted with mammalian cells and it was determined that crucially, the films remain highly biocompatible.

**Figure 1:**
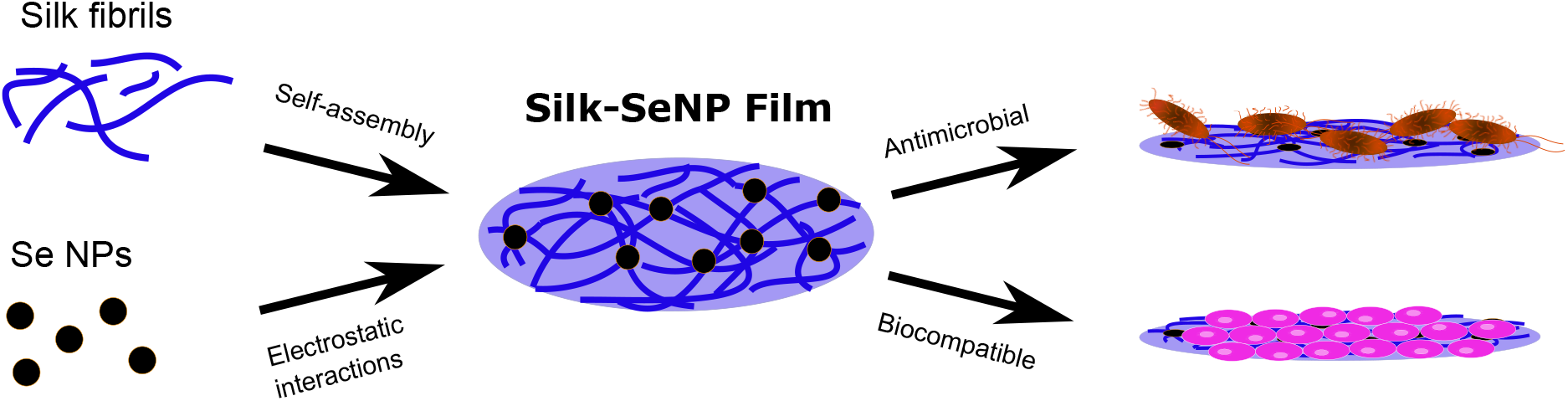
Schematic showing the formation of the Silk-SeNP films and their use as antimicrobial agents.

As described above, selenium nanoparticles (SeNPs) were first synthesised. A range of sodium selenite concentrations were investigated, while the ascorbic acid solution, which was in excess with respect to the sodium selenite, was kept constant at a concentration of 56.7 mM. Following the reduction reaction, the solution was left at room temperature for 24 hours in order to promote SeNP formation. Selenium nanoparticles were imaged by transmission electron microscopy (TEM) and scanning electron microscopy (SEM). From the TEM micrographs, it was determined that the average sizes of the nanoparticles for the 7 different concentrations of sodium selenite investigated, ranged from around 60 to 130 nm. Moreover, depending on the concentration of sodium selenite, we could systematically control the nanoparticle size. The largest SeNPs were formed using a sodium selenite concentration of 200 *μ*g/mL.

For each sodium selenite concentration used, the size of more than 100 nanoparticles was measured. The mean size (*μ*) and the standard deviation (*σ*) of the nanoparticles was thus determined. The average coefficient of variation (which is the ratio of *σ* to *μ* and gives an indication of the dispersion of the distribution) for these samples was calculated as 9.8%, demonstrating a fairly narrow distribution for most of the nanoparticle systems synthesised. The average sizes of the SeNPs for the 7 conditions tested can be seen in Figure 2a, while some representative TEM and SEM micrographs of the nanoparticles can be seen in Figures 2b-c and 2d-e, respectively.

**Figure 2:**
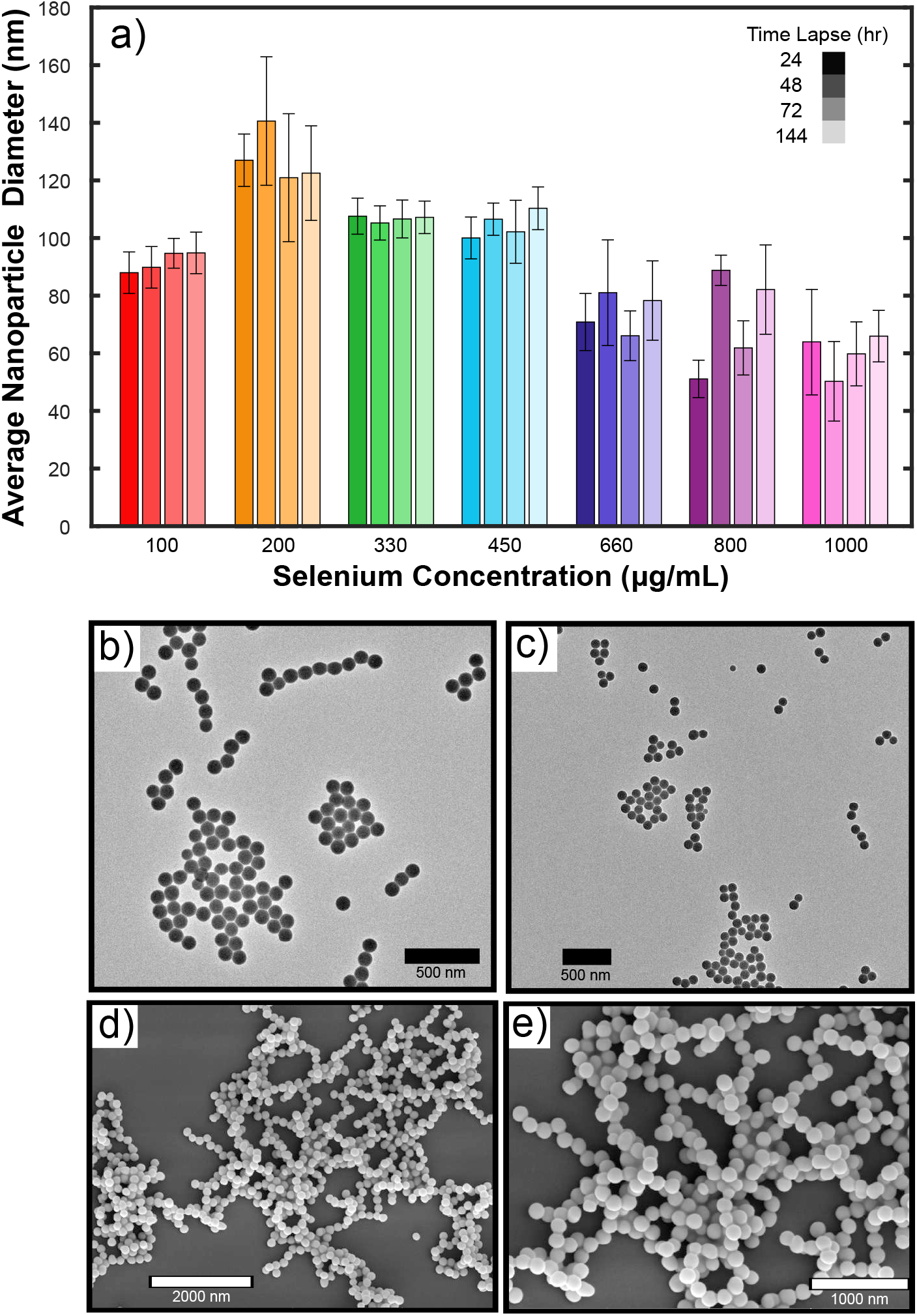
**(a)** Average diameter of SeNPs at 24 hr, 48 hr, 72 hr, and 144 hr as a function of selenium nanoparticle concentration for seven samples: (red) 100 *μ*g/mL, (orange) 200 *μ*g/mL, (green) 330 *μ*g/mL, (light blue) 450 *μ*g/mL, (dark blue) 660 *μ*g/mL, (purple) 800 *μ*g/mL, and (pink) 1000 *μ*g/mL. TEM micrographs for two samples: **(b)** 200 *μ*g/mL and **(c)** 800 *μ*g/mL. **(d-e)** SEM micrographs of SeNPs at a concentration of 200 *μ*g/mL.

Next, we explored the stability of the SeNPs over time. It is known that with time, nanoparticles have the tendency to degrade and form aggregated structures, resulting in changes in their size distribution. In order to monitor this process, we looked at the size distribution of the SeNPs for all 7 systems over a period of 144 hours. TEM was employed to monitor the time dependent size distribution and again, more than 100 nanoparticles were measured for each bar graph, as shown in Figure 2a. It was determined that for all the systems, SeNPs remained stable over time with minimal size discrepancies. The different colours in Figure 2a represent the different nanoparticle concentrations, while the colour gradient corresponds to the day on which the measurements were made. It is clear that the size distribution of the SeNPs follows the same trend in day 2 as well as in day 6.

Moreover, in order to probe the elemental composition of the nanoparticles, energy-dispersive X-ray spectroscopy (EDX) was performed on a nanoparticle sample. As can be seen in SI Figure S2, the Se peaks at 1.5 keV confirm that the nanoparticles imaged were in fact selenium-based, with other peaks characteristic of the copper grid used for sample deposition and of the staining agent, uranyl acetate. Furthermore, powder X-ray diffraction (PXRD) was conducted on a 600 *μ*g/mL nanoparticle solution. The peak positions (shown in red) are representative of a pure hexagonal phase of selenium crystals with lattice parameters a = 4.366 and c = 4.9536^44^ (SI Figure S3).

To monitor the stability of the nanoparticles, the use of UVvisible absorption spectroscopy was employed. Absorption spectra of SeNPs were taken and are depicted in Figures 3a-b. The absorption intensity of the nanoparticles at their characteristic wavelength (265 nm) is an indication of the stability, where the higher the intensity, the more stable the nanoparticles.^43^ From the absorption spectra in Figure 3a, which correspond to the different sodium selenite concentrations tested, it can be seen that the most stable system is at 1000 *μ*g/mL, while the least stable nanoparticle system is at 330 *μ*g/mL. Moreover, by monitoring an intermediate stability nanoparticle system over time, at 200 *μ*g/mL, it was determined that there is a clear decrease in the absorbance at 265 nm (Figure 3b), which indicates that the nanoparticles start to clump and aggregate over time.

**Figure 3:**
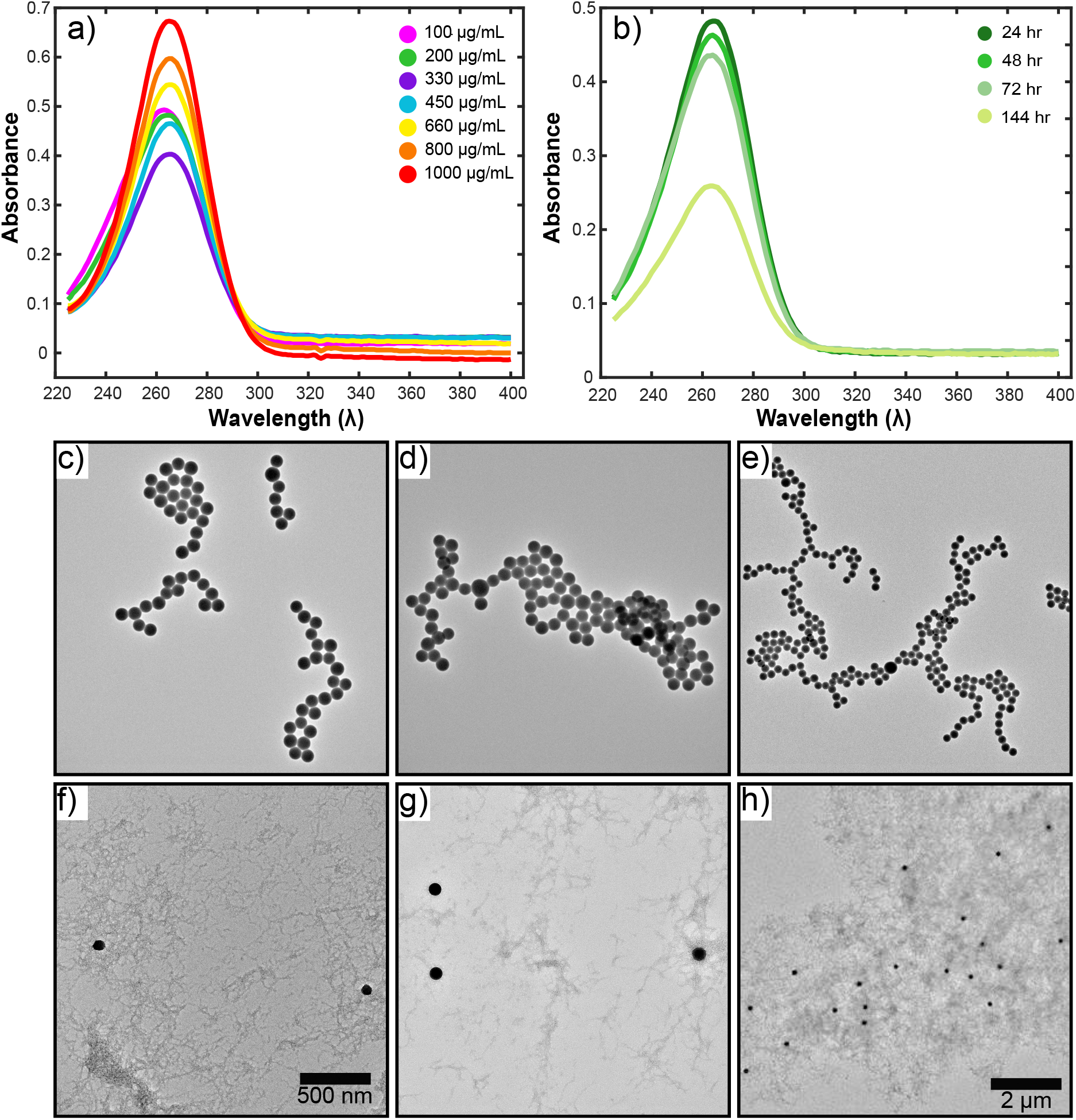
**(a)** UVvisible absorption spectra for SeNPs taken at day 2. Seven different samples were tested: (pink) 100 *μ*g/mL, (green) 200 *μ*g/mL, (purple) 330 *μ*g/mL, (blue) 450 *μ*g/mL, (yellow) 660 *μ*g/mL, (orange) 800 *μ*g/mL and (red) 1000 *μ*g/mL. **(b)** UVvisible absorption spectra for a 200 *μ*g/mL nanoparticle solution at 24 hr, 48 hr, 72 hr, and 144 hr. **(c-e)** TEM micrographs of SeNP’s without silk fibrils and **(f-h)** with silk fibrils. Silk fibrils prevent the nanoparticles from agglomerating.

It was determined that the nanoparticles aggregated for all concentrations tested. Moreover, even though this agglomeration process was initiated from day 3 onwards, complete agglomeration occurred by day 6, as can be seen in the TEM micrographs in Figures 3c-e. This was also corroborated from the absorbance spectra at 265 nm (Figure 3b), where it is clear that the intensity of the nanoparticle spectra decreases over time. Furthermore, massive clumps, on the order of millimetres, were seen after 2 weeks of formation (SI Figure S1). In order to address the issue of nanoparticle agglomeration, regenerated silk fibroin (RSF) was used. RSF is an FDA approved protein, which has the propensity to self-assemble into nanofibrils. By combining SeNPs with silk nanofibrils, we were able to stabilise the nanoparticles and prevent agglomeration. The nanoparticles had the tendency to adsorb to the fibrillar network and were therefore spatially immobilised. This mechanism of stabilisation was confirmed via TEM, and some typical micrographs can be seen in Figures 3f-h.

Having established that silk fibrils stabilise SeNPs against aggregation, we next sought to form a material capable of exhibiting antimicrobial properties. The silk fibril-SeNP (SF-SeNP) hybrid inorganic/organic solution was mixed with a 40% ethanol solution (which is a strong gel promoter) at a 1:1 ratio, and the final mixture was left to dry at room temperature for one day. The experimental procedure used to make the films is shown in the schematic in Figure 4a. This process resulted in the formation of a SF-SeNP film (as can be seen in Figure 4 and in SI Figures S4b-c). By allowing the solution to dry at room temperature, a film with a uniform nanoparticle distribution could be formed (Figure 4). In order to monitor the antimicrobial properties of the films, we prepared SF-SeNP films for the seven different concentrations tested, i.e. the 50 500 *μ*g/mL concentration range. The films were then cut using a hole puncher and placed in a 96-well plate, in order to conduct absorbance and fluorescence related assays using a plate-reader experiment.

**Figure 4:**
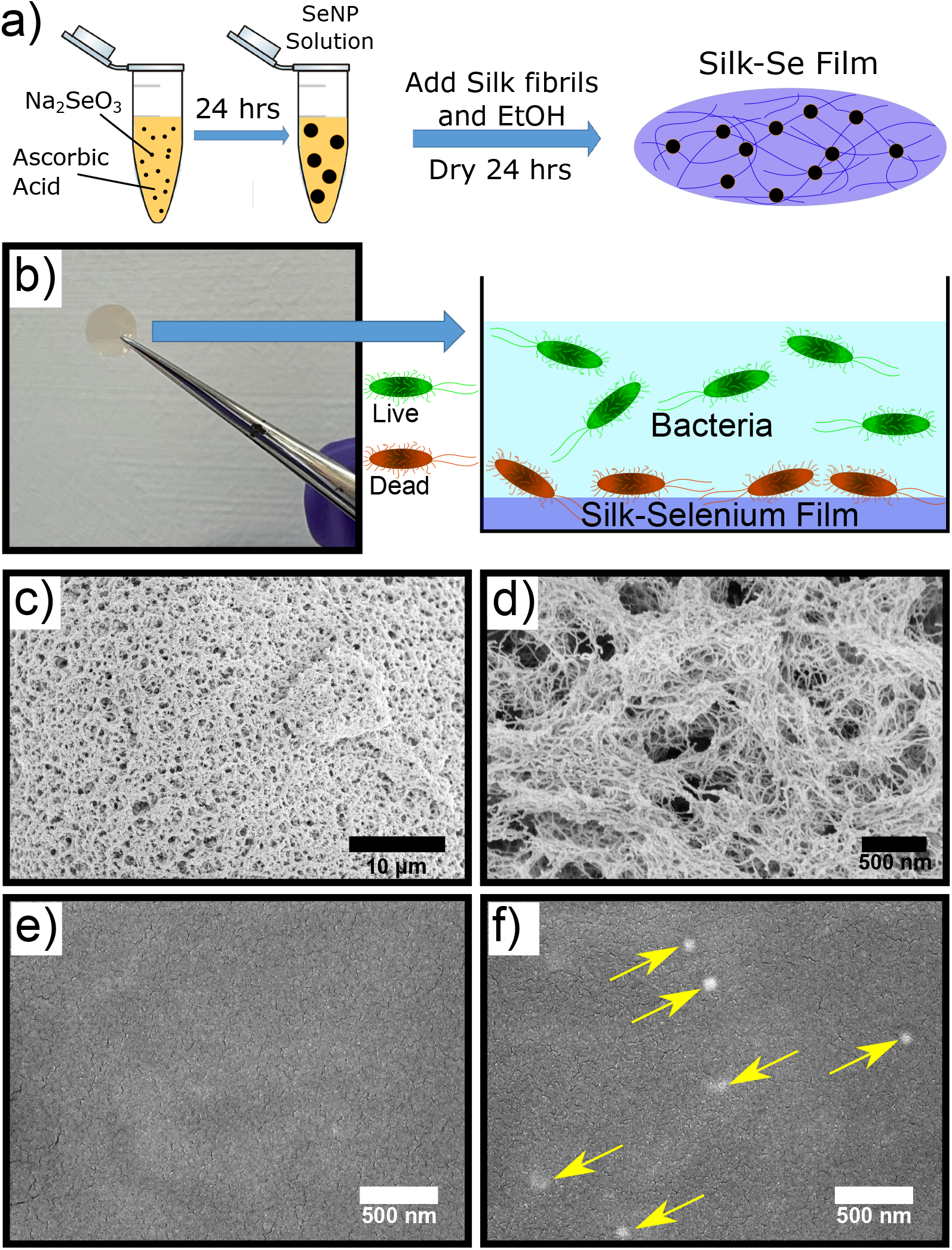
**(a-b)** Schematic representation of the experimental procedure used to make the silk-selenium films and the experimental setup in which antimicrobial inhibition was monitored. **(c-d)** High magnification SEM micrographs of silk films showing the highly dense fibrillar network. **(e-f)** SEM micrographs of silk films in: **(d)** the absence of selenium nanoparticles and **(e)** the presence of selenium nanoparticles. The yellow arrows point towards the nanoparticles within the film.

Before investigating the antimicrobial properties of the SF-SeNP films, we first characterised the films using scanning electron microscopy (SEM). Micrographs of both silk films with and without selenium nanoparticles were taken. High magnification SEM micrographs revealed the highly dense fibrillar network with the silk films (Figures 4c-d). A representative image of a silk film without nanoparticles can be seen in Figure 4e. It contains a homogeneous distribution of the self-assembled protein network. Moreover, in the micrographs of the SF-SeNP films (Figure 4f), the nanoparticles can clearly be seen, with yellow arrows pointing towards the selenium nanoparticles within the film.

Following film formation, we then evaluated the antibacterial activity of the SF-SeNP films against *E. coli*. Direct assessment of bacterial viability of *E. coli* was carried out using live/dead viability analysis. This was achieved through use of Syto9 (a dye that stains and indicates live bacteria) and propidium iodide (a dye that stains and indicates dead bacteria). Following 24h of treatment, bacterial cell death was observed for samples treated with the SF-SeNP films (bottom panel of figure 5a). This effect was observed for a range of nanoparticle concentrations, with the 500 *μ*g/mL system showing the highest potency. Furthermore, the control and silk film treated bacteria remained viable and proliferated over time, as shown in Figure 5a (top and middle panels, respectively).

**Figure 5:**
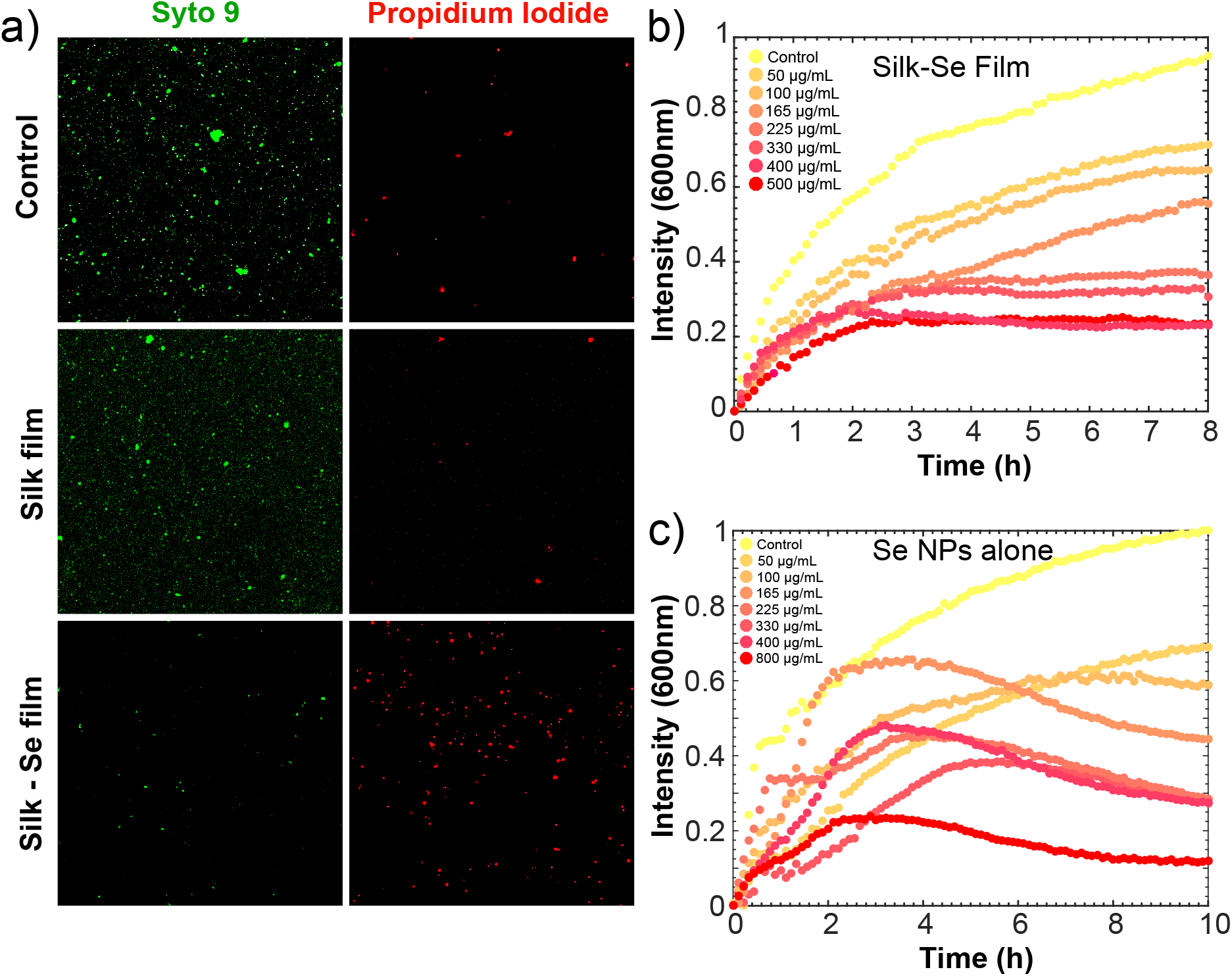
**(a)** Bacterial viability analysis of *E. coli* via live/dead staining following overnight treatment with and without silk-selenium films. A 40X magnification was used for all micrographs. **(b)** Kinetics of bacterial growth inhibition on a silk-selenium film via turbidity analysis. **(c)** Kinetics of bacterial growth inhibition using bare selenium nanoparticles obtained via turbidity analysis. The absorbance measurements were done at 600 nm.

Moreover, the effect of selenium nanoparticle concentration on bacterial cell death was investigated. This was done by conducting a kinetic growth inhibition analysis. *E. coli* cultures, grown to an OD of 0.2, were added to the nanoparticle containing films and to the corresponding controls. The bacteria were incubated overnight and absorbance measurements at 600 nm were conducted over time. It is evident from Figure 5b that there is a dose-dependent trend, with the higher the nanoparticle concentration, the more potent the antibacterial properties of the film. Treatment with the silk films alone did not affect bacterial growth (Figure 5b). Moreover, it was observed that the SF-SeNP films with concentrations of 225 *μ*g/mL and above, had >70% bacterial death.

In order to assess whether the SF-SeNP films retain their antibacterial potency as opposed to using just bare nanoparticles, the kinetic inhibition analysis was also performed for free nanoparticles in solution. The bacteria were again incubated overnight and absorbance measurements at 600 nm were conducted over time. It is evident from SI Figure 5c that a similar dose-dependent trend is observed, as in the case of the SF-SeNP films. It was found that with bare nanoparticles, a similar amount of bacteria were killed (~70% bacterial death) for nanoparticle concentrations up to 400 *μ*g/mL. Therefore we can conclude that our SF-SeNP films retain their antibacterial action and have similar performances to using free nanoparticles.

Furthermore, the antibacterial activity of the films on a gram-positive bacterium, *B. subtilis*, was investigated. It has been shown that due to the selenium nanoparticles having an overall negative charge, they are more potent on gram-positive bacteria. ^45^ It was therefore expected that for the same nanoparticle concentrations, the antibacterial activity of the films would be more potent on *B. subtilis* than on the gram-negative bacterium *E. coli*. Bacterial viability was again assessed using Syto9 and propidium iodide. Following 24h of treatment, bacterial cell death was observed for samples treated with the SF-SeNP films (bottom panel of figure 6a), whereas the control and silk film treated bacteria remained both viable and proliferated over time (top and middle panels of Figure 6a, respectively).

**Figure 6:**
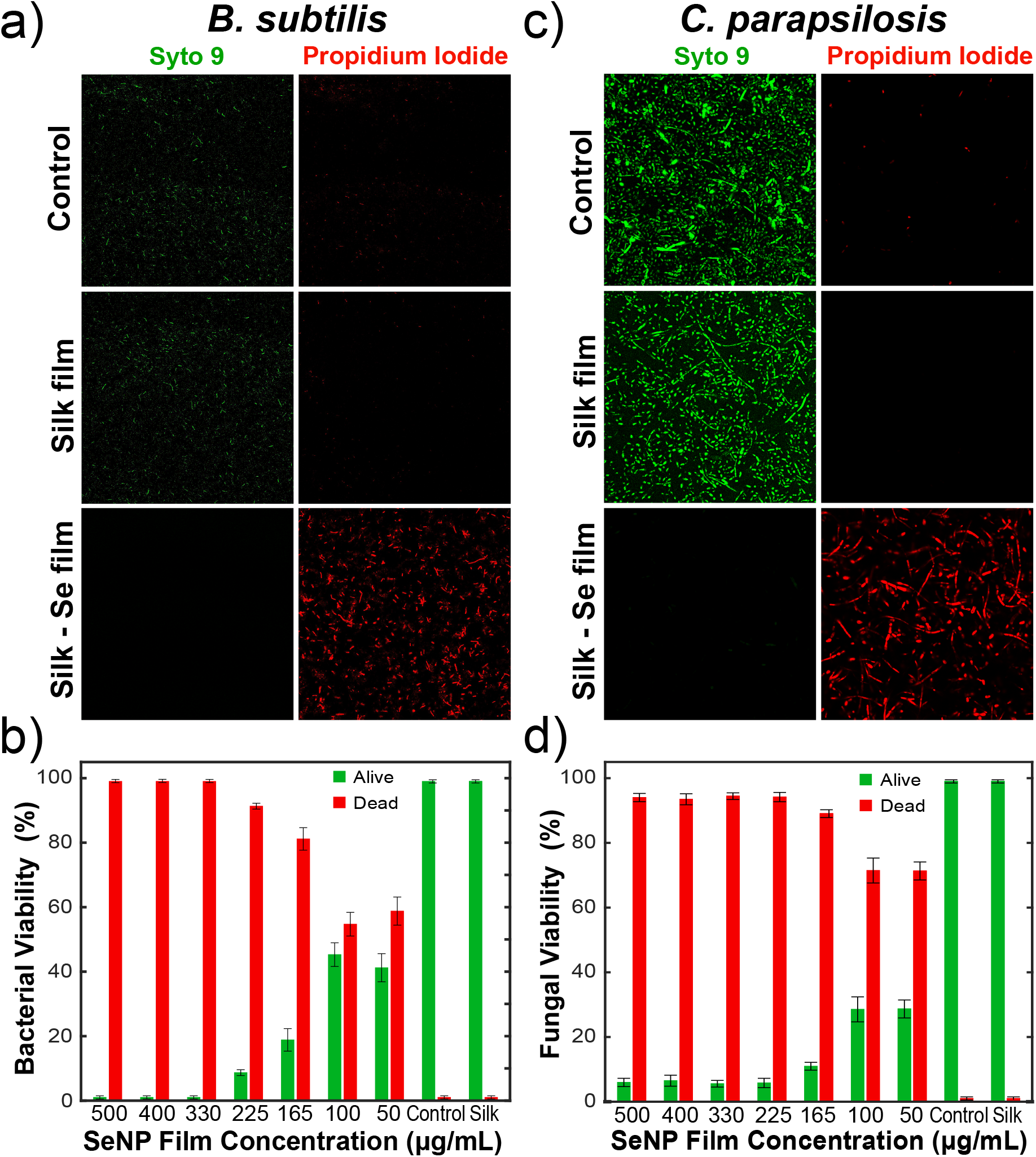
**(a)** Bacterial viability analysis of *B. subtilis* via live/dead staining following overnight treatment. A 40X magnification was used for all micrographs. **(b)** Quantification of bacterial viability of *B. subtilis* on selenium films. **(c)** Fungal viability analysis of *C. parapsilosis* via live/dead staining following overnight treatment. A 40X magnification was used for all micrographs. **(d)** Quantification of fungal viability of *C. parapsilosis*on selenium films.

We next quantified the viability of the *B. subtilis* as a function of nanoparticle concentration within the films, as shown in Figure 6b. This was done by counting the live/dead cells from the confocal images taken. For each film condition, an average of 10 images was used, which resulted in hundreds of cells being counted per condition. Our results indicate that at a concentration of 300 *μ*g/mL and above, total bacterial eradication was observed, whereas at a concentration of 225 *μ*g/mL, more than 92% of *B. subtilis* was killed. It can be seen in the live/dead viability analysis that even at really low nanoparticle concentrations (50 *μ*g/mL), ca. 60% bacterial death was observed.

Moreover, the antifungal activity of the films was investigated on the fungi *Candida parapsilosis*. Previous literature has shown that selenium nanoparticles have good antifungal activity. ^40^ We found that our hybrid films had a very potent antifungal effect. Fungal viability was assessed using Syto9 and propidium iodide. Samples were again incubated for a total of 24 h at 30°C. It was observed that for samples treated with the SF-SeNP films (bottom panels of figure 6c) total fungal death was observed, whereas the control and silk film treated fungi remained viable and also proliferated over time (top and middle panels of figure 6c, respectively). We quantified the effect of the films on *C. parapsilosis* by counting the live/dead cells from the confocal images, using the same approach as previously mentioned (Figure 6d). The antifungal effect of the SF-SeNP films is more potent than their antibacterial effect, with around 73% fungal death observed for the films with nanoparticle concentrations as low as 50 *μ*g/mL. More importantly, the concentration which kills the majority of fungi (>95%) is 225 *μ*g/mL and above.

In order to gain a mechanistic insight into the mode of action of the SF-SeNP films, a membrane permeation assay was conducted with SYTOX Blue. SYTOX Blue is a cationic dye that cannot enter an intact cell, but rather it can only enter if the membrane has been disrupted. The dye binds to intracellular nucleic acids and is fluorescent when excited at 405 nm. ^20^ Samples treated with SF-SeNP films had significantly higher fluorescence signal compared to control samples, indicating that membrane disruption was evident, and is thus the probable mode of action behind the antimicrobial effect of the SF-SeNP films (SI Figure S5).

Finally, in order to assess the ability of these films as agents for treating topical infections, the biocompatibility of the antibacterial films with mammalian cells (HEK-293) via an MTT-based cell viability assay was evaluated. HEK-293 cells were grown in 96-well plates overnight in the presence of the films and controls. Cell viability was not affected by the presence of a silk film. Moreover, in the presence of films containing low nanoparticle concentrations (100 *μ*g/mL and below), high cellular viability was observed (Figure 7b). Cellular viability was reduced to just over 85% when treated with films containing a Se NP concentration of 225 *μ*g/mL. However, the viability was reduced to around 20% when the cells were treated with films containing 400 and 500 *μ*g/mL (Figure 7b).

**Figure 7:**
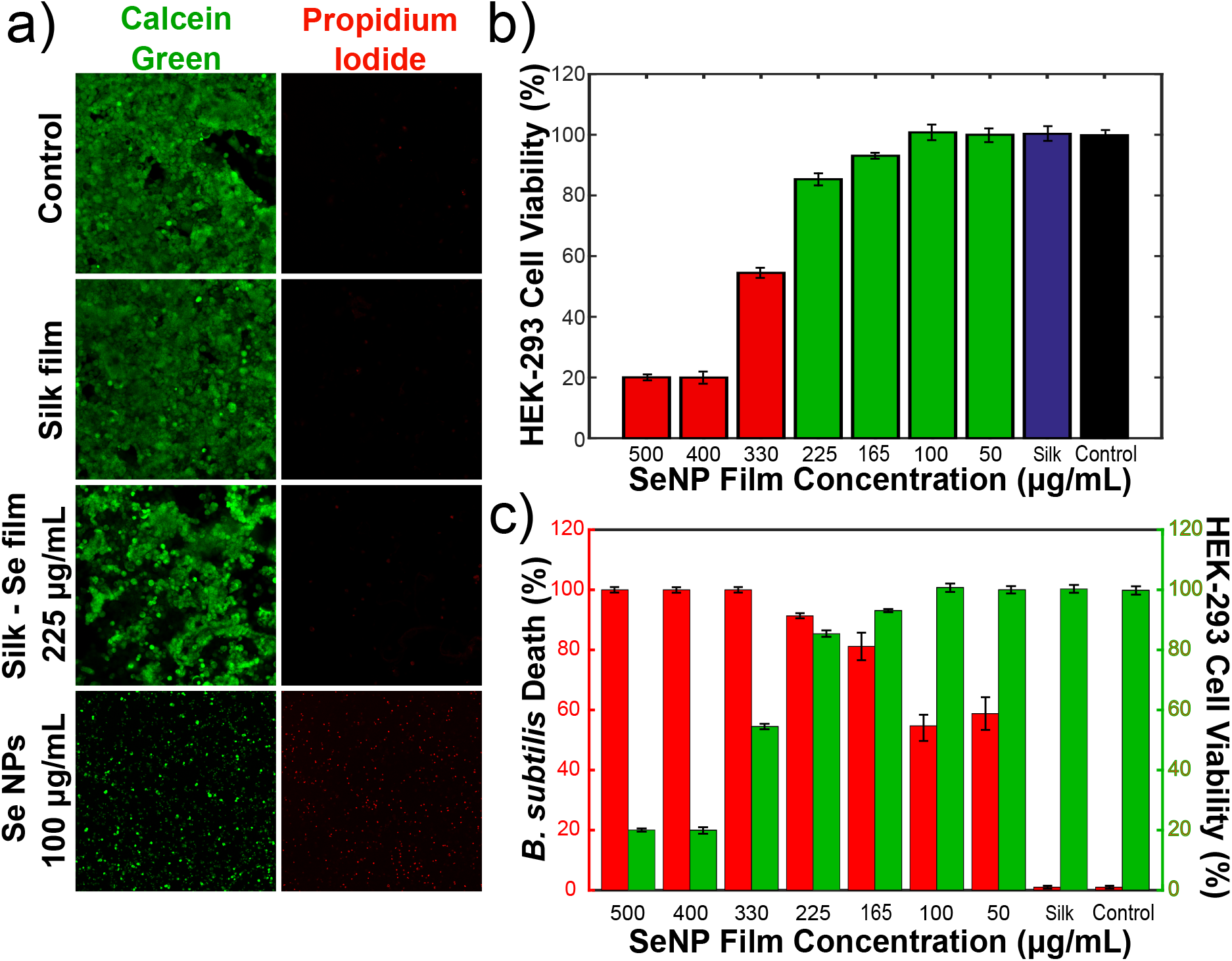
**(a)** Mammalian cell viability analysis with the antibacterial films and controls following an overnight incubation. This was conducted using a fluorescence-based live/dead staining assay containing fluorescein diacetate (live cells) and propidium iodide (dead cells). A 40X magnification was used for all the microscopy images. **(b)** MTT cell viability analysis of HEK-293 cells with the antibacterial films and controls. **(c)** Combined graph of the antibacterial activity on *B. subtilis* and HEK-293 cell viability as a function of silk-selenium film concentration.

Additionally, a live/dead analysis of HEK-293 cells which were treated in a similar manner indicated the same results. Calcein AM staining (indicating live cells) and ethidium homodimer-1 (indicating dead cells) was conducted (Figure 7a). From the images, it is clear that there is minimal cell death in the presence of films, both with and without nanoparticles (top three panels of figure 7a). However, in the presence of free nanoparticles, cell viability is greatly reduced, with high cellular death and inhibition of normal cell proliferation (bottom panel of figure 7a). Our results thus show that by encapsulating the nanoparticles within these silk films, mammalian cells can be shielded from the cytotoxic effects, making them more biocompatible, whilst also providing high antimicrobial activity.

Taken together, the biocompatability results and the antimicrobial results strongly suggest that an optimum concentration of SF-SeNP films exists. Using a concentration of 225 *μ*g/mL we were able to eradicate the majority of bacteria (>70% for *E. coli* and >92% for *B. subtilis*) and fungi (>95%), whilst also protecting and keeping the majority of HEK-293 cells alive (>85%). This is evident in Figure 7c, which shows the graph of *B. subtilis* death (in red) and HEK-293 cell viability (in green) as a function of silk-selenium film concentration.

## Conclusion

Antimicrobial resistance is an increasingly major healthcare problem of increasing urgency. It has become clear that alternative routes, in addition to conventional antibiotic and antifungal treatments, need to be developed. In this study we have combined selenium nanoparticles with silk fibroin in order to form biocompatible films that exhibit strong antimicrobial action. Two different bacteria, gram-negative *E. coli* and grampositive *B. subtilis* were tested, while the fungi *C. parapsilosis* was also investigated. In all cases, it was found that the films displayed potent antibacterial and antifungal properties. Furthermore, the biocomatibility of the films was investigated via an MTT-based cell viability assay. HEK-293 mammalian cell viability was monitored and it was established that in the presence of films containing low nanoparticle concentrations, high cellular viability was observed. This is primarily because the protein scaffold has the ability to shield and protect the mammalian cells from the cytotoxic effects of the bare nanoparticles. By combining the results from the antimicrobial assays and the mammalian cell viability assays, it was determined that the optimum concentration which displays both high biocompatability and high bacterial and fungal death was 225 *μ*g/mL. This optimum film condition had the ability to kill the majority of bacteria and fungi whilst also keeping the majority of HEK-293 cells alive. As bacteria and fungi are unlikely to develop antimicrobial resistance to the nanoparticles embedded within the films, these hybrid organic/inorganic bioinspired materials provide a basis for potential biomedical and biotechnological applications, especially for use as topical antimicrobial agents.

## Acknowledgements

We would like to thank Walther C. Traberg for his assistance with cell culture and Prof. Roisin Owens for kindly providing the HEK293 cells. We would like to acknowledge Heather Greer for her assistance in acquiring TEM images and EDX spectra of the Se nanoparticles presented in this paper. We would also like to acknowledge Samuel Dada and Akhila J. Denduluri for their helpful comments concerning confocal microscopy and viability assays. We would like to acknowledge the EPSRC Underpinning Multi-User Equipment Call (EP/P030467/1) for funding the TEM in the Yusuf Hamied Department of Chemistry. We would also like to acknowledge the UKRI World Class Laboratories (WCL) fund for funding the Leica TCS SP8 inverted confocal microscope used for imaging in this paper. E.G.W. is funded by the Gates Cambridge Trust. A.K.J acknowledges funding from the Cambridge Trust, the EPSRC grant EP/L015978/1 for the Centre for Doctoral Training for Nanoscience and Nanotechnology (NanoDTC), Queens’ College. T.P.J.K. acknowledges funding from the European Research Council under the European Unions Seventh Framework Programme (FP7/2007-2013) through the ERC grants PhysProt (agreement no. 337969), the Biotechnology and Biological Sciences Research Council (BBSRC), the Frances and Augustus Newman Foundation, and the Centre for Misfolding Diseases.

## Materials and Methods

### Formation and Purification of Regenerated Silk Fibroin

Regenerated silk fibroin was obtained from Bombyx mori silkworm cocoons, following a previously determined protocol.^15^ In summary, cut cocoons were boiled for 30 minutes in a solution of 0.02 M sodium carbonate. The remaining silk product was rinsed with cold water and dried overnight. A 1:4 ratio of 9.3 M lithium bromide solution was added to the silk fibroin and heated at 60 C for 4 h. The salt was then removed using dialysis and the product was incubated for 2 days with several water changes. The resulting silk fibroin solution was purified by centrifugation and stored at 4 C.

### Synthesis of Selenium Nanoparticles

Selenium nanoparticles were prepared by optimizing a previously established protocol in which sodium selenite is reduced by ascorbic acid.^43^ Briefly, 5 samples of 1800 L Na2O3Se in Milli-Q water were prepared (100 *μ*g/mL, 200 *μ*g/mL, 330 *μ*g/mL, 450 *μ*g/mL, 660 *μ*g/mL, 800 *μ*g/mL, and 1000 *μ*g/mL). Ascorbic acid (200 L, 56.7 mM) was added dropwise to the sodium selenite solution and was mixed thoroughly to obtain the selenium nanoparticles. A color change occurred upon nanoparticle formation from translucent colorless to translucent red.

### Characterization of Selenium Nanoparticles

TEM, SEM, UVvisible absorption spectroscopy, DLS, and PXRD were carried out to characterize our products. The size and distribution of the selenium nanoparticles were studied by transmission electron microscopy (TEM). TEM micrographs were recorded using a Thermo Scientific (FEI) Talos F200X G2 electron microscope. Samples were prepared over a period of six days in which nanoparticles were drop-casted on carbon-coated TEM copper grids and dried after 30 seconds of exposure. The resulting TEM micrographs were analyzed using Siemens Totally Integrated Automation (TIA) software and ImageJ software.

Scanning Electron Microscopy was employed to image the nanoparticles using a FEI Verios 460 SEM at a current of 5 kV. The selenium nanoparticle solution was mounted onto a silicon wafer by adding 1 *μ*L of solution and letting the sample air dry. This wafer was placed onto a multipin specimen holder, and a 10 nm platinum layer was sputter coated onto the sample.

Selenium nanoparticles were also characterized by UVvisible absorption spectroscopy (Implen NanoPhotometer NP80). The samples were diluted by a factor of 100 and the spectra were recorded within a 200400 nm wavelength range over a period of 6 days. The average particle size (hydrodynamic diameter) of the synthesized nanoparticles was measured using dynamic light scattering (DLS). The samples were diluted by a factor of 1000 and the spectra were recorded on a Malvern Zetasizer Nano instrument. Nanoparticle composition was characterized using a Bruker D8-QUEST PHOTON-100 diffractometer. The resulting crystallite size was compared to the standard crystalline selenium diffraction pattern.

### Incorporation of Selenium Nanoparticles into Silk Fibroin Films

A solution of regenerated silk fibroin was mixed in a 1:1 ratio with ethanol. This was in turn mixed with a 1:1 ratio with each concentration of selenium nanoparticles, yielding final nanoparticle concentrations of 50, 100, 165, 225, 300, 400 and 500 *μ*g/mL. The final solutions were pipetted onto a weighing boat and left to dry overnight, resulting in the formation of a film. The film was peeled off and cut using a hole puncher. This was put into a 96-well plate in order to conduct the antimicrobial assays.

### PXRD Analysis

Powder X-ray diffraction (PXRD) of the 660 *μ*g/mL of selenium nanoparticle sample was carried out using a Panalytical Empyrean diffractometer. The sample was added dropwise in 5 applications to a single crystal silicon sample holder and dried at 50 C between applications. The sample was measured for 15 hours. The selenium peaks in the spectra match the expected reference peaks indicated by the red lines.

### Transmission electron microscopy (TEM) and energy-dispersive X-ray (EDX) spectroscopy analysis

Transmission electron microscopy (TEM), high angle annular dark field scanning transmission electron microscopy (HAADF-STEM) and energy-dispersive X-ray (EDX) spectroscopy were performed using a Thermo Scientific (FEI) Talos F200X G2 TEM operating at 200 kV. TEM images were acquired using a Ceta 16M CMOS camera. EDX spectra were collected using the Super-X EDS detector system which consists of 4 windowless silicon drift detectors. The peak positions in SI Figure S3 (shown in red) recorded at 2*θ* = 23.9, 30.0, 41.7, 44.0, 45.7, 52.0, 56.4, 62.2, 65.5 and 68.4 are characteristic of the (100), (101), (110), (102), (111), (201), (003), (202), (210) and (211) respectively, and are representative of a pure hexagonal phase of selenium crystals with lattice parameters a = 4.366 and c = 4.9536^44^ (SI Figure S3). TEM grids (continuous carbon film on 300 mesh Cu) were glow discharged using a Quorum Technologies GloQube instrument at a current of 25mA for 60s. Samples were either observed as prepared or negatively stained using a 2% uranyl acetate solution for 45s.

### Kinetic Growth Analysis

*E. coli* bacterial cells were grown to an OD600 of 0.2 in LB media. Film samples were placed within the wells of a 96-well plate and bacterial solutions were added on top. Kinetic growth inhibition was determined by turbidity analysis via optical density measurements (at 600 nm) using a FLUOstar Omega microplate reader (BMG Labtech). Three independent experiments were conducted and the results presented are indicative of the triplicates.

### Viability analysis of bacterial and fungal cells

*E. coli* and *C. parapsilosis* bacteria were incubated at 37°C with films containing nanoparticles of various concentrations in well chambers. Similarly, *C. parapsilosis* fungi was incubated at 30°C with films containing nanoparticles in the same format. Post incubation, bacteria were extracted using a pipette and transferred to an eppendorf tube. Syto 9 and propidium iodide (LIVE/DEAD BacLight Bacterial Viability Kit, Thermo Fisher Scientific, TFS) were subsequently added in a 1:1 ratio following a 10x dilution. The bacterial cells were immediately observed under a 40x oil objective using a Leica TCS SP8 inverted confocal microscope.

### Confocal Microscopy and Cell Membrane Study

Confocal microscopy was employed to study the mechanism behind bacteria eradication. SYTOX Blue (Thermo Fisher Scientific) 1 *μ*M was incubated with *E. coli* at an OD600 of 0.1 for 30 minutes at 37°C. The sample was then examined using confocal microscopy LSM 510, excited at 405 nm (Leica TCS SP8).

### Cell culture of HEK-293 cells

Human embryonic kidney 293 (HEK-293) cells were cultured in 25 cm^2^ flasks at 37°C and 5% CO_2_ using advanced Dulbeccos modified Eagle medium (DMEM; TFS) with the addition of 10% fetal bovine serum (Merck), 50 U/mL penicillin and 50 *μ*g/mL streptomycin (TFS), 1% (v/v) GlutaMax (TFS), and 50 *μ*g/mL gentamicin (TFS).

### Cytotoxicity and cell proliferation using MTT Assay on HEK-293 cells

The viability of HEK-293 cells following incubation with films was determined using a standard MTT cell proliferation assay. Approximately 10^5^ cells were seeded per well in a 96 well plate and incubated for 24h under 37°C and 5% CO_2_. Following this, 10 L of MTT (3-[4,5-dimethylthiazol-2-yl]-2,5-diphenyltetrazolium bromide) labelling reagent Merck) was added to each well with an incubation period of 4 hours. Post-incubation, 100 L of the solubilisation solution was added to each well and a further overnight incubation was performed at 37°C and 5% CO_2_. The absorbance of the resulting purple solution was measured at 595 nm using a FLUOstar Omega microplate reader (BMG Labtech).

### Viability analysis of HEK-293 cells

The viability of HEK-293 cells post exposure to films was also determined using a LIVE/DEAD Viability/Cytotoxicity Kit (Invitrogen). Approximately 10^5^ cells were seeded per well, in which films were pre-deposited. The cells were incubated for 24 hours, and a stock solution containing 5 L calcein AM (Component A) and 20 L ethidium homodimer-1 (Component B) was added to 10 mL Dulbecco’s PBS to create a stock solution. 100-200 L of the stock dye solution was added to each well and the cells were observed under 10x and 40x objectives in a Leica TCS SP8 inverted confocal microscope to determine the relative number of live and dead cells.

## Supporting Information

**Figure S1:**
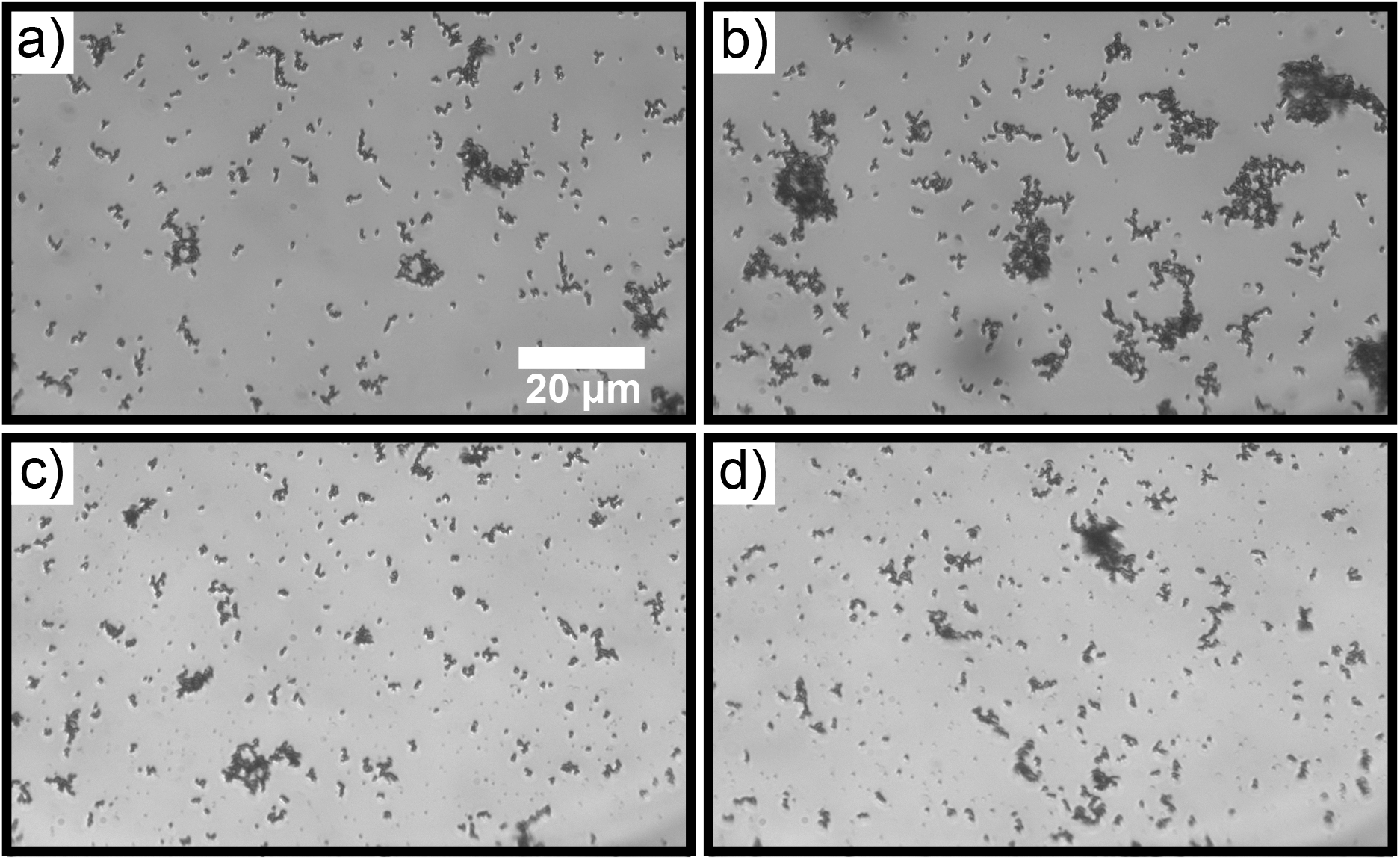
Se nanoparticle agglomeration monitored over time. After day 6 of formation, it was observed that the nanoparticles tend to clump into micrometer-sized structures.

**Figure S2:**
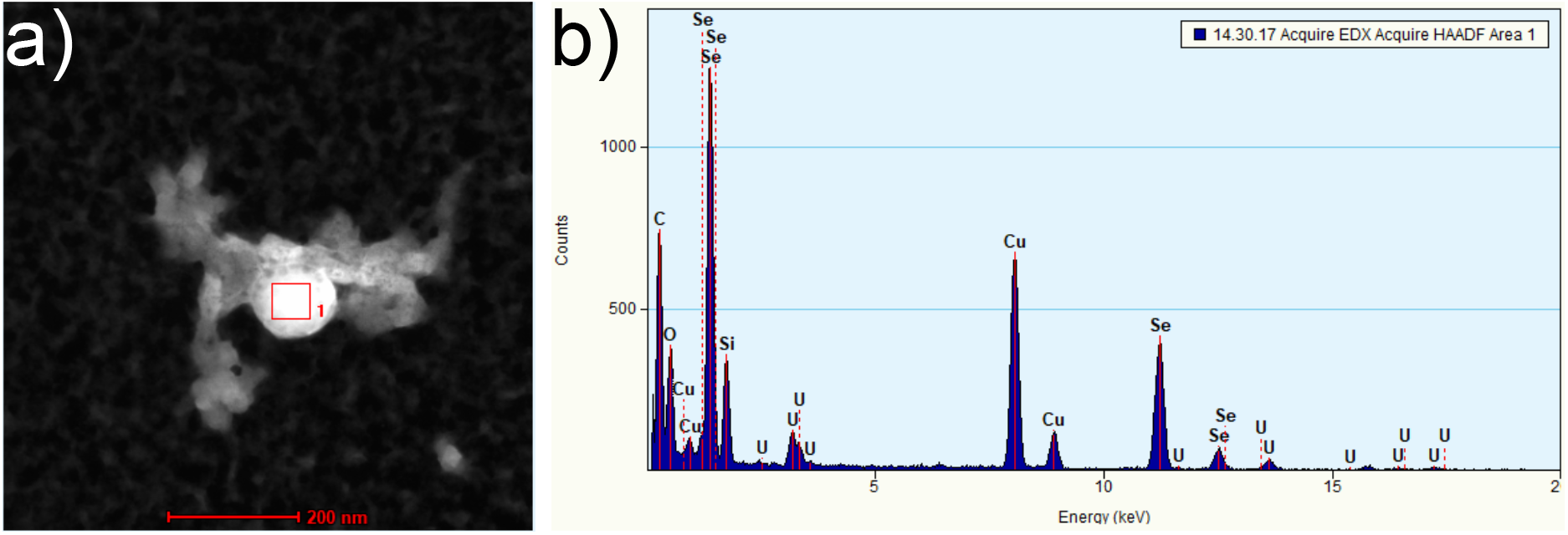
EDX spectrum of a 600 *μ*g/mL selenium nanoparticle solution.

**Figure S3:**
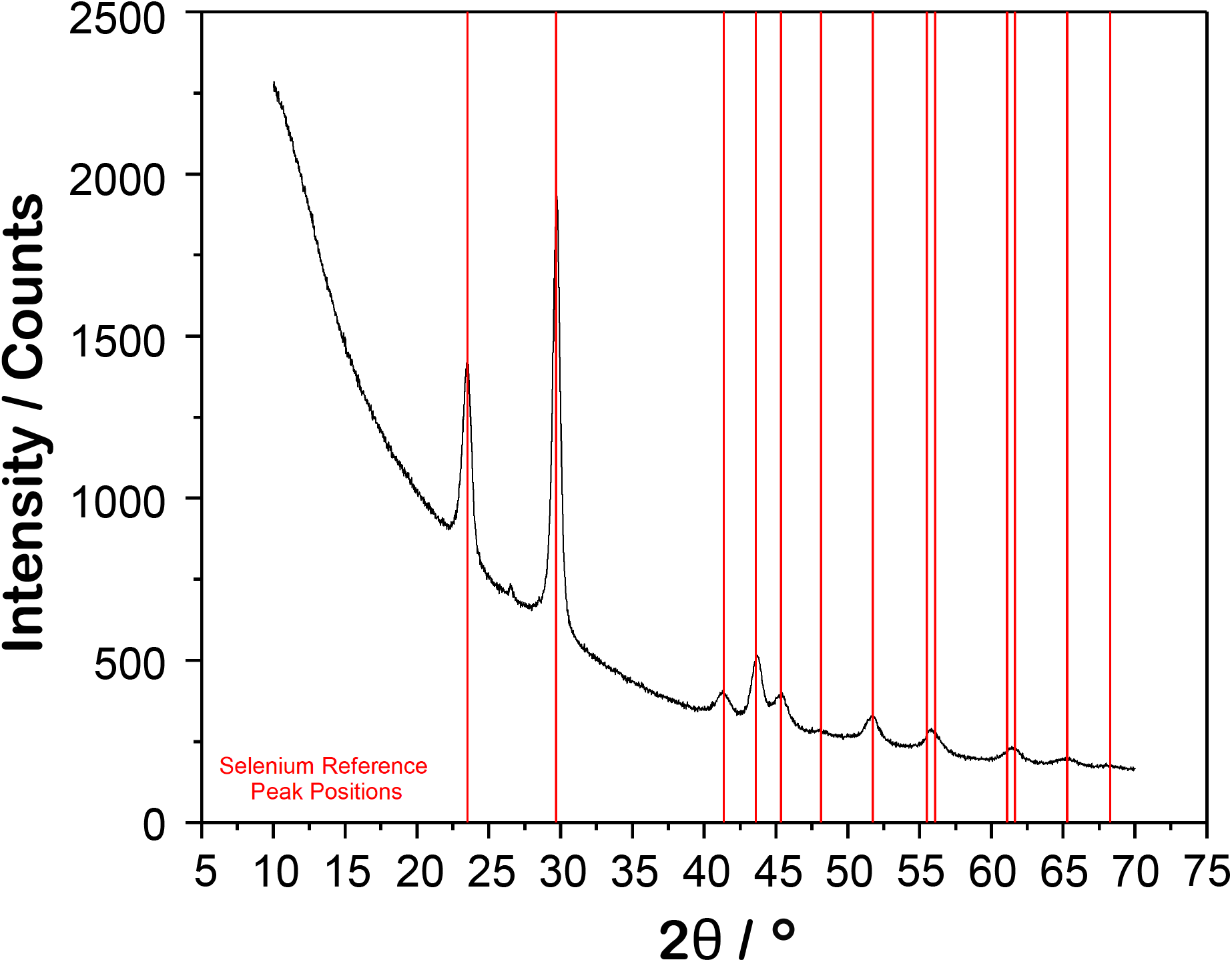
XRD spectrum of a 600 *μ*g/mL selenium nanoparticle solution.

**Figure S4:**
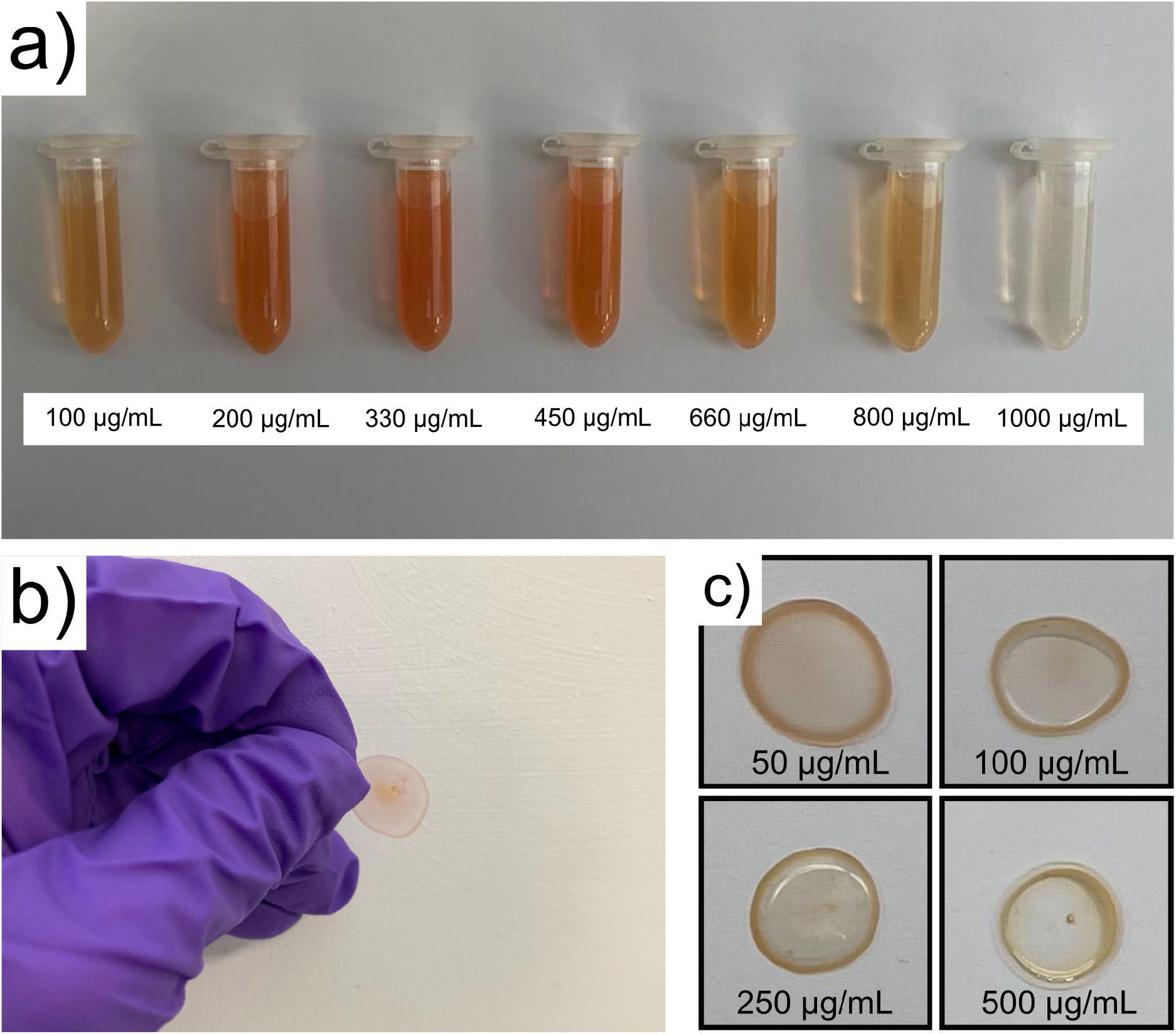
(**a**) Image of the 7 SeNP systems tested following 1 day of incubation. (**b-c**) Images of films made using the hybrid inorganic/organic silk SeNP system.

**Figure S5:**
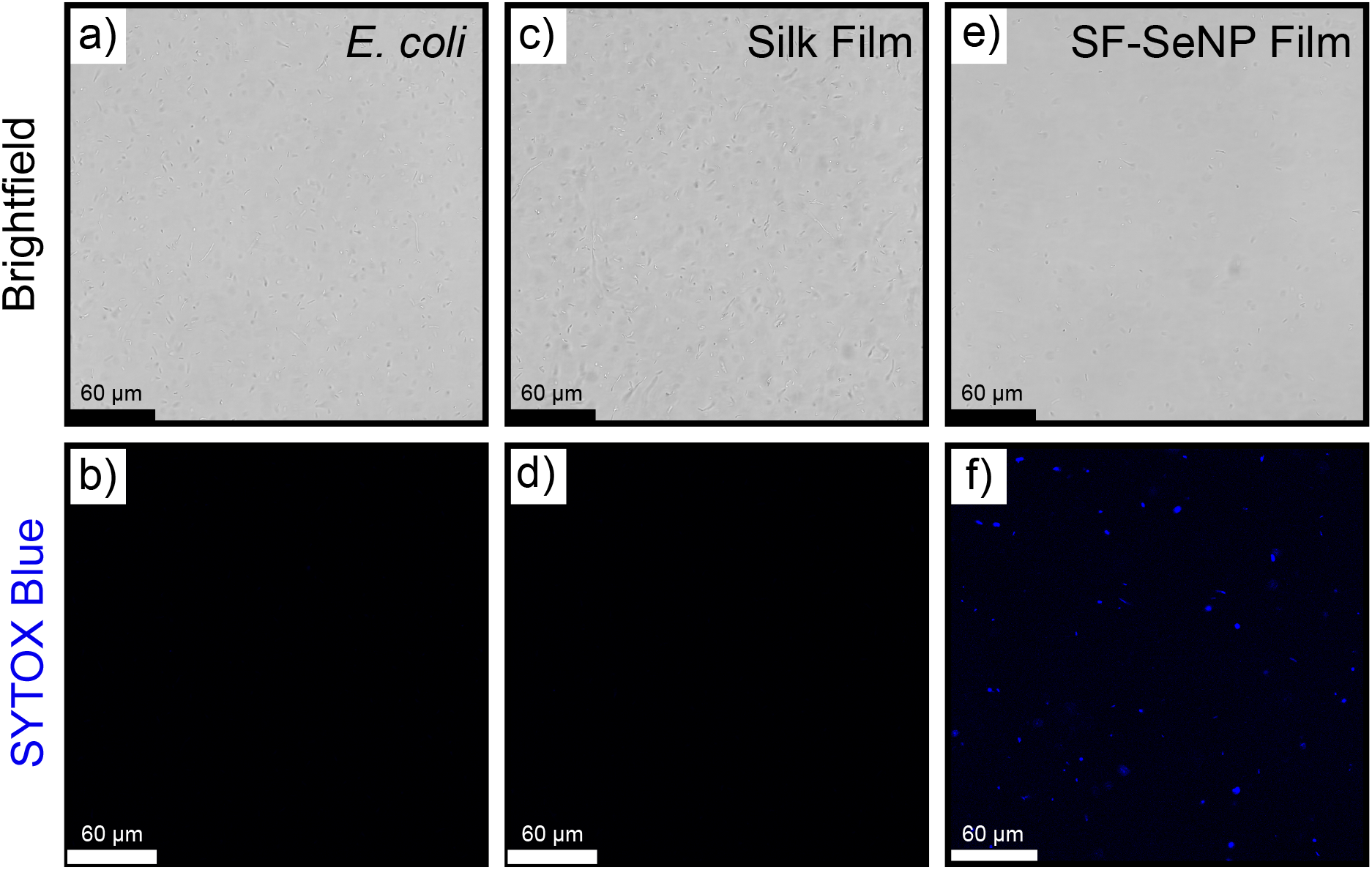
(**a-f**) Brightfield and confocal micrographs of *E. coli* with and without the presence of silk films. (**a-b**) *E. coli* control. (**c-d**) *E. coli* incubated with silk films. Brightfield images indicate that bacteria proliferated on the film while no SYTOX Blue staining of cell membranes was observed (**e-f**) *E. coli* incubated with SF-SeNP Films. Brightfield images show a decreased number of bacteria. SYTOX Blue staining of bacterial membranes was observed, indicating membrane disruption.

## References

(1) Zhang, S. Fabrication of novel biomaterials through molecular self-assembly. Nature Biotechnology 2003, 21, 1171–1178.

(2) Schnaider, L.; Toprakcioglu, Z.; Ezra, A.; Liu, X.; Bychenko, D.; Levin, A.; Gazit, E.; Knowles, T. P. Biocompatible Hybrid Organic/Inorganic Microhydrogels Promote Bacterial Adherence and Eradication in Vitro and in Vivo. Nano Letters 2020, 20, 1590–1597.

(3) Shen, Y.; Levin, A.; Kamada, A.; Toprakcioglu, Z.; Rodriguez-Garcia, M.; Xu, Y.; Knowles, T. P. From Protein Building Blocks to Functional Materials. ACS Nano 2021, 15, 5819–5837.

(4) Yuan, C.; Ji, W.; Xing, R.; Li, J.; Gazit, E.; Yan, X. Hierarchically oriented organization in supramolecular peptide crystals. Nature Reviews Chemistry 2019, 3, 567–588.

(5) Adler-Abramovich, L.; Gazit, E. The physical properties of supramolecular peptide assemblies: From building block association to technological applications. Chemical Society Reviews 2014, 43, 6881–6893.

(6) Shimanovich, U.; Michaels, T. C.; De Genst, E.; Matak-Vinkovic, D.; Dobson, C. M.; Knowles, T. P. Sequential Release of Proteins from Structured Multishell Microcapsules. Biomacromolecules 2017, 18, 3052–3059.

(7) Varanko, A.; Saha, S.; Chilkoti, A. Recent trends in protein and peptide-based biomaterials for advanced drug delivery. Advanced Drug Delivery Reviews 2020, 156, 133–187.

(8) Liu, X.; Toprakcioglu, Z.; Dear, A. J.; Levin, A.; Ruggeri, F. S.; Taylor, C. G.; Hu, M.; Kumita, J. R.; Andreasen, M.; Dobson, C. M.; Shimanovich, U.; Knowles, T. P. Fabrication and Characterization of Reconstituted Silk Microgels for the Storage and Release of Small Molecules. Macromolecular Rapid Communications 2019, 40.

(9) Khalid, A.; Mitropoulos, A. N.; Marelli, B.; Tomljenovic-Hanic, S.; Omenetto, F. G. Doxorubicin loaded nanodiamond-silk spheres for fluorescence tracking and controlled drug release. Biomedical Optics Express 2016, 7, 132.

(10) Hakala, T. A.; Davies, S.; Toprakcioglu, Z.; Bernardim, B.; Bernardes, G. J.; Knowles, T. P. A Microfluidic Co-Flow Route for Human Serum Albumin-DrugNanoparticle Assembly. Chemistry - A European Journal 2020, 26, 5965–5969.

(11) Toprakcioglu, Z.; Knowles, T. P. Shear-mediated sol-gel transition of regenerated silk allows the formation of Janus-like microgels. Scientific Reports 2021, 11.

(12) Toprakcioglu, Z.; Kumar Challa, P.; Morse, D. B.; Knowles, T. Attoliter protein nanogels from droplet nanofluidics for intracellular delivery. Science Advances 2020, 6.

(13) Xing, R.; Liu, K.; Jiao, T.; Zhang, N.; Ma, K.; Zhang, R.; Zou, Q.; Ma, G.; Yan, X. An Injectable Self-Assembling Collagen-Gold Hybrid Hydrogel for Combinatorial Antitumor Photothermal/Photodynamic Therapy. Advanced Materials 2016, 28, 3669–3676.

(14) Koh, L. D.; Cheng, Y.; Teng, C. P.; Khin, Y. W.; Loh, X. J.; Tee, S. Y.; Low, M.; Ye, E.; Yu, H. D.; Zhang, Y. W.; Han, M. Y. Structures, mechanical properties and applications of silk fibroin materials. Progress in Polymer Science 2015, 46, 86–110.

(15) Rockwood, D. N.; Preda, R. C.; Yücel, T.; Wang, X.; Lovett, M. L.; Kaplan, D. L. Materials fabrication from Bombyx mori silk fibroin. Nature Protocols 2011, 6, 1612–1631.

(16) Guo, C.; Li, C.; Vu, H. V.; Hanna, P.; Lechtig, A.; Qiu, Y.; Mu, X.; Ling, S.; Nazarian, A.; Lin, S. J.; Kaplan, D. L. Thermoplastic moulding of regenerated silk. Nature Materials 2020, 19, 102–108.

(17) Rizzo, G.; Lo Presti, M.; Giannini, C.; Sibillano, T.; Milella, A.; Guidetti, G.; Musio, R.; Omenetto, F. G.; Farinola, G. M. Bombyx mori Silk Fibroin Regeneration in Solution of Lanthanide Ions: A Systematic Investigation. Frontiers in Bioengineering and Biotechnology 2021, 9.

(18) Vollrath, F.; Porter, D. Spider silk as archetypal protein elastomer. Soft Matter 2006, 2, 377–385.

(19) Magaz, A.; Roberts, A. D.; Faraji, S.; Nascimento, T. R.; Medeiros, E. S.; Zhang, W.; Greenhalgh, R. D.; Mautner, A.; Li, X.; Blaker, J. J. Porous, Aligned, and Biomimetic Fibers of Regenerated Silk Fibroin Produced by Solution Blow Spinning. Biomacromolecules 2018, 19, 4542–4553.

(20) Schnaider, L.; Ghosh, M.; Bychenko, D.; Grigoriants, I.; Ya’Ari, S.; Shalev Antsel, T.; Matalon, S.; Sarig, R.; Brosh, T.; Pilo, R.; Gazit, E.; Adler-Abramovich, L. Enhanced Nanoassembly-Incorporated Antibacterial Composite Materials. ACS Applied Materials and Interfaces 2019,

(21) Chung, S.; Webster, T. J. Electrospun Silk with Selenium Nanoparticles Inhibits Bacterial Proliferation. 2016,

(22) Zou, X.; Jiang, Z.; Li, L.; Huang, Z. Selenium nanoparticles coated with pH responsive silk fibroin complex for fingolimod release and enhanced targeting in thyroid cancer. Artificial Cells, Nanomedicine and Biotechnology 2021, 49, 83–95.

(23) Larsson, D. G.; Flach, C. F. Antibiotic resistance in the environment. 2021.

(24) Prestinaci, F.; Pezzotti, P.; Pantosti, A. Antimicrobial resistance: A global multifaceted phenomenon. Pathogens and Global Health 2015, 109, 309–318.

(25) Debroas, D.; Siguret, C. Viruses as key reservoirs of antibiotic resistance genes in the environment. ISME Journal 2019, 13, 2856–2867.

(26) AshaRani, P. V.; Mun, G. L. K.; Hande, M. P.; Valiyaveettil, S. Cytotoxicity and genotoxicity of silver nanoparticles in human cells. ACS Nano 2009, 3, 279–290.

(27) Le Ouay, B.; Stellacci, F. Antibacterial activity of silver nanoparticles: A surface science insight. Nano Today 2015, 10, 339–354.

(28) Salas-Orozco, M.; Niño-Martínez, N.; Martínez-Castañón, G. A.; Méndez, F. T.; Jasso, M. E. C.; Ruiz, F. Mechanisms of resistance to silver nanoparticles in endodontic bacteria: A literature review. 2019.

(29) Rayman, M. P. Selenium and human health. Lancet 2012, 379, 1256–68.

(30) Karnaukh, E. A.; Walker, L. M.; Lynch, K. A.; Wiita, E. G.; Buzzeo, M. C. Electrochemical Study of Selenocystine Reactivity and Reduction at Metallic Surfaces. ChemElectroChem 2017, 4, 1250–1255.

(31) Evans, S. O.; Khairuddin, P. F.; Jameson, M. B. Optimising selenium for modulation of cancer treatments. Anticancer Research 2017, 37, 6497–6509.

(32) Filipović, N.; Ušjak, D.; Milenković, M. T.; Zheng, K.; Liverani, L.; Boccac-cini, A. R.; Stevanovic, M. M. Comparative Study of the Antimicrobial Activity of Selenium Nanoparticles With Different Surface Chemistry and Structure. Frontiers in Bioengineering and Biotechnology 2021, 8.

(33) Kuršvietienė, L.; Mongirdienė, A.; Bernatonienė, J.; Šulinskienė, J.; Stanevišienė, I. Selenium anticancer properties and impact on cellular redox status. Antioxidants 2020, 9.

(34) Chen, W.; Li, Y.; Yang, S.; Yue, L.; Jiang, Q.; Xia, W. Synthesis and antioxidant properties of chitosan and carboxymethyl chitosan-stabilized selenium nanoparticles. Carbohydrate Polymers 2015, 132, 574–581.

(35) Zou, J.; Su, S.; Chen, Z.; Liang, F.; Zeng, Y.; Cen, W.; Zhang, X.; Xia, Y.; Huang, D. Hyaluronic acid-modified selenium nanoparticles for enhancing the therapeutic efficacy of paclitaxel in lung cancer therapy. Artificial Cells, Nanomedicine and Biotechnology 2019, 47, 3456–3464.

(36) Mary, T. A.; Shanthi, K.; Vimala, K.; Soundarapandian, K. PEG functionalized selenium nanoparticles as a carrier of crocin to achieve anticancer synergism. RSC Advances 2016, 6, 22936–22949.

(37) Qiu, W. Y.; Wang, Y. Y.; Wang, M.; Yan, J. K. Construction, stability, and enhanced antioxidant activity of pectin-decorated selenium nanoparticles. Colloids and Surfaces B: Biointerfaces 2018, 170, 692–700.

(38) Pi, J.; Jiang, J.; Cai, H.; Yang, F.; Jin, H.; Yang, P.; Cai, J.; Chen, Z. W. Ge11 peptide conjugated selenium nanoparticles for egfr targeted oridonin delivery to achieve enhanced anticancer efficacy by inhibiting egfr-mediated pi3k/akt and ras/raf/mek/erk pathways. Drug Delivery 2017, 24, 1549–1564.

(39) Ferro, C.; Florindo, H. F.; Santos, H. A. Selenium Nanoparticles for Biomedical Applications: From Development and Characterization to Therapeutics. 2021.

(40) Geoffrion, L. D.; Hesabizadeh, T.; Medina-Cruz, D.; Kusper, M.; Taylor, P.; Vernet-Crua, A.; Chen, J.; Ajo, A.; Webster, T. J.; Guisbiers, G. Naked Selenium Nanoparticles for Antibacterial and Anticancer Treatments. ACS Omega 2020,

(41) Guisbiers, G.; Wang, Q.; Khachatryan, E.; Mimun, L. C.; Mendoza-Cruz, R.; Larese-Casanova, P.; Webster, T. J.; Nash, K. L. Inhibition of e. Coli and s. aureus with selenium nanoparticles synthesized by pulsed laser ablation in deionized water. International Journal of Nanomedicine 2016, 11, 3731–3736.

(42) Lara, H. H.; Guisbiers, G.; Mendoza, J.; Mimun, L. C.; Vincent, B. A.; Lopez-Ribot, J. L.; Nash, K. L. Synergistic antifungal effect of chitosan-stabilized selenium nanoparticles synthesized by pulsed laser ablation in liquids against Candida albicans biofilms. International Journal of Nanomedicine 2018, 13, 2697–2708.

(43) Vahdati, M.; Tohidi Moghadam, T. Synthesis and Characterization of Selenium Nanoparticles-Lysozyme Nanohybrid System with Synergistic Antibacterial Properties. Scientific Reports 2020, 10.

(44) Cruz, L. Y.; Wang, D.; Liu, J. Biosynthesis of selenium nanoparticles, characterization and X-ray induced radiotherapy for the treatment of lung cancer with interstitial lung disease. Journal of Photochemistry and Photobiology B: Biology 2019, 191, 123–127.

(45) Arakha, M.; Pal, S.; Samantarrai, D.; Panigrahi, T. K.; Mallick, B. C.; Pramanik, K.; Mallick, B.; Jha, S. Antimicrobial activity of iron oxide nanoparticle upon modulation of nanoparticle-bacteria interface. Scientific Reports 2015, 5.

